# Emergence of XDR high-risk *Pseudomonas aeruginosa* ST309 in South America: a global comparative genomic analysis

**DOI:** 10.1101/2021.01.21.427610

**Authors:** Érica L. Fonseca, Sérgio M. Morgado, Raquel V. Caldart, Fernanda Freitas, Ana Carolina P. Vicente

**Affiliations:** Laboratório de Genética Molecular de Microrganismos, Instituto Oswaldo Cruz, FIOCRUZ, RJ, Brazil; Universidade Federal de Roraima, Boa Vista, Roraima, Brazil

**Keywords:** High-risk clone, pandemic lineage, ST309, Brazil, mobilome, heavy metal resistance

## Abstract

*Pseudomonas aeruginosa* has been considered one of the major nosocomial pathogens associated with elevated morbidity and mortality worldwide. Outbreaks have been associated with few high-risk pandemic *P. aeruginosa* lineages, presenting a remarkable antimicrobial resistance. However, the biological features involved with the persistence and spread of such lineages among clinical settings remain to be unravel. This study reports the emergence of the ST309 *P. aeruginosa* lineage in South America/Brazil, more precisely, in the Amazon region. Global genomic analyses were performed with the Brazilian strain (PA834) and more 41 complete and draft ST309 genomes publicly available, giving insights about ST309 epidemiology and its resistome and mobilome. Antimicrobial susceptibility tests revealed that the Brazilian PA834 strain presented the XDR phenotype, which was mainly due to intrinsic resistance mechanisms. Genomic analyses revealed a heterogeneous distribution of acquired antimicrobial resistance genes among ST309 genomes, which included *bla*_VIM-2_, *bla*_IMP-15_ and *qnrVC1*, all of them associated with class 1 integrons. The mobilome mining showed the presence of Integrative and Conjugative Elements, transposons and genomic islands harbouring a huge arsenal of hevy metal resistance genes. Moreover, these elements also carried genes involved with virulence and adaptive traits. Therefore, the presence of such genes in ST309 lineage possibly accounted for the global spread and persistence of this emerging clone, and for its establishment as a pandemic lineage of clinical importance.

## 1. Introduction

The opportunistic pathogen *Pseudomonas aeruginosa* has been considered one of the major nosocomial pathogens associated with elevated morbidity and mortality worldwide [1]. Outbreaks have been associated with high-risk pandemic *P. aeruginosa* lineages, which present a remarkable antimicrobial resistance due to a notable multifactorial antimicrobial resistance mechanism, which leads to the emergence of multidrug (MDR) and extensively drug resistance (XDR) phenotypes [2]. In addition, some *P. aeruginosa* lineages harbour mechanisms of metals and organic solvent resistance, which contribute to their persistence in hospital settings and pandemic potencial [2–4].

The *P. aeruginosa* ST309 lineage has been recently assigned as a potential emerging threat [5], since it has already been involved with outbreaks in Asia (Korea), Europe (Greece and France), North and Central Americas (USA and Mexico) [5–9]. Therefore, studies focusing on ST309 mobilome would bring insights on the evolution of this lineage, concerning the virulence, resistance and metabolic features that contribute to its success in spreading and establishing infections worldwide. In this study we reported the emergence of ST309 in South America (PA834), and performed a global genomic epidemiologic analyses that revealed ST309 as a high-risk pandemic lineage, probably due to a mobilome carrying a remarkable arsenal of heavy metal resistance, virulence and antimicrobial resistance genes (ARGs).

## 2. Materials and methods

### 2.1. Bacterial strain, Antimicrobial Susceptibility Test and MLST analysis

In 2016, a carbapenem-resistant *P. aeruginosa* (PA834) was recovered from a blood infection case, from a 65-year old male placed in the medical clinic ward of the General Hospital of Roraima, placed in Boa vista, a city embedded in the Amazon region. This patient remained hopitalized for 8 months, and was treated with imipenem, ceftazidime and ciprofloxacin prior bacterium isolation. Species identification was carried out by the automated VITEK2 System.

The antimicrobial susceptibility test was determined by Disc-diffusion method (bioMérieux, France) on Mueller-Hinton medium for the following antibiotics: gentamicin, amikacin, tobramycin, kanamycin, neomycin, streptomycin, spectinomycin, aztreonam, cefalotin, cefoxitin, cefuroxime, ceftazidime, ceftriaxone, cefotaxime, cefepime, ampicillin/sulbactam, amoxacillin, amoxacillin/clavulanate, penicillin, ticarcillin/clavulanate, piperacillin, piperacillin/tazobactam, oxacillin, azitromycin, claritromycin, erithromycin, nalidixic acid, ciprofloxacin, norfloxacin, levofloxacin, ofloxacin, sulfonamide, trimethoprim, trimethoptim/sulfamethoxazole, tetracycline, clindamycin, fosfomycin, lincomycin, nitrofurantoine, rifampin, tigecycline. The MIC of carbapenems (imipenem, meropenem, ertapenem and doripenem) was determined by the E-test method (bioMérieux, France). The MICs of polymyxin B and colistin were assessed by the broth microdilution method with antibiotic concentrations ranged from 0.1 μg/mL to 64 μg/mL (MIC breakpoint for resistance > 2 μg/mL, according to CLSI/EUCAST guidelines) [10]. The current definition criterium for classifying *P. aeruginosa* antimicrobial resistance was applied [11]. The carbapenemase production was verified by the modified Hodge Test [12].

### 2.2. Whole Genome Sequencing and genome annotation

PA834 genomic DNA was obtained by using the NucleoSpin Microbial DNA kit (Macherey-Nagel), and genomic library was prepared using Nextera XT paired-end run with a ~500-bp insert. The whole genome sequencing was performed with the Illumina HiSeq 2500 sequencer at the High-Throughput Sequencing Platform of the Oswaldo Cruz Foundation (Fiocruz, Rio de Janeiro, Brazil). The quality of the reads was assessed with FASTQC and *de novo* assembling was obtained with the SPAdes 3.5 assembler with default settings. Gene prediction and annotation were performed with Rapid Annotations using Subsystems Technology (RAST) server and Prokka software (https://github.com/tseemann/prokka).

The finished genome assembly and annotated sequences for PA834 were deposited in GenBank under accession number VRZB00000000, with Bioproject number PRJNA560372 (URLs - https://www.ncbi.nlm.nih.gov/nuccore/1780198729).

### 2.3. Comparative genomic analyses and in silico characterization of the resistome, mobilome and virulence genes

All complete and draft *P. aeuruginosa* genomes (n=5,344) (last updated on August, 2020) were retrieved from GenBank database. The seven MLST alleles that define the ST309 (*acsA-13, aroE-8, guaA-9, mutL-3, nuoD-1, ppsA-17*, and *trpE-15*) were separetely used as query in BLASTn analyses to recover all currently available ST309 genomes. Additionally, some ST309 genomes were also recovered from EMBL database based on information provided in the Bacterial Isolate Genome Sequence Database (BIGSdb). A total of 41 ST309 genomes, recovered worldwide in distinct spatio-temporal contexts, were used in comparative genomic analyses together with the Brazilian PA834 genome (Table 1).

**Table 1.**
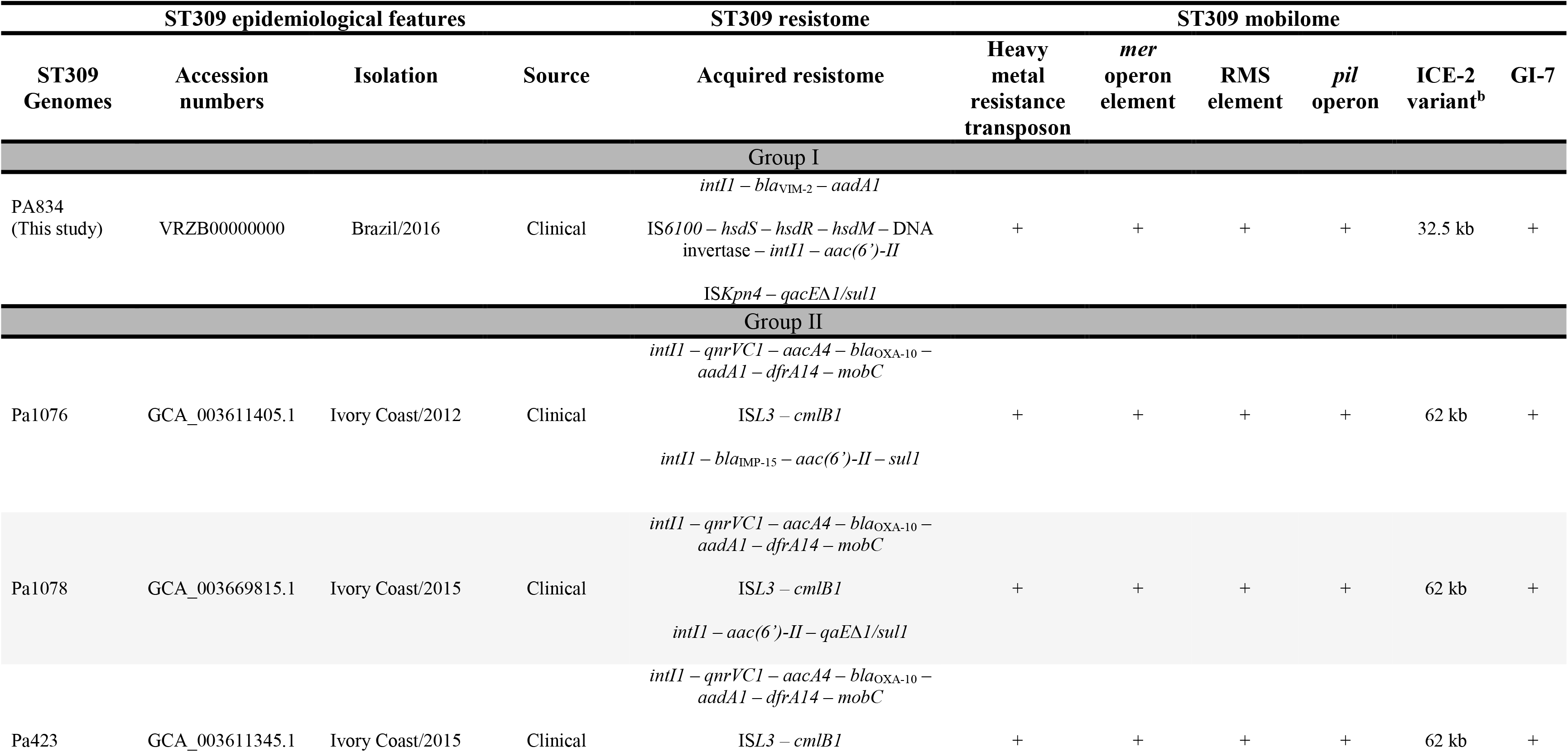
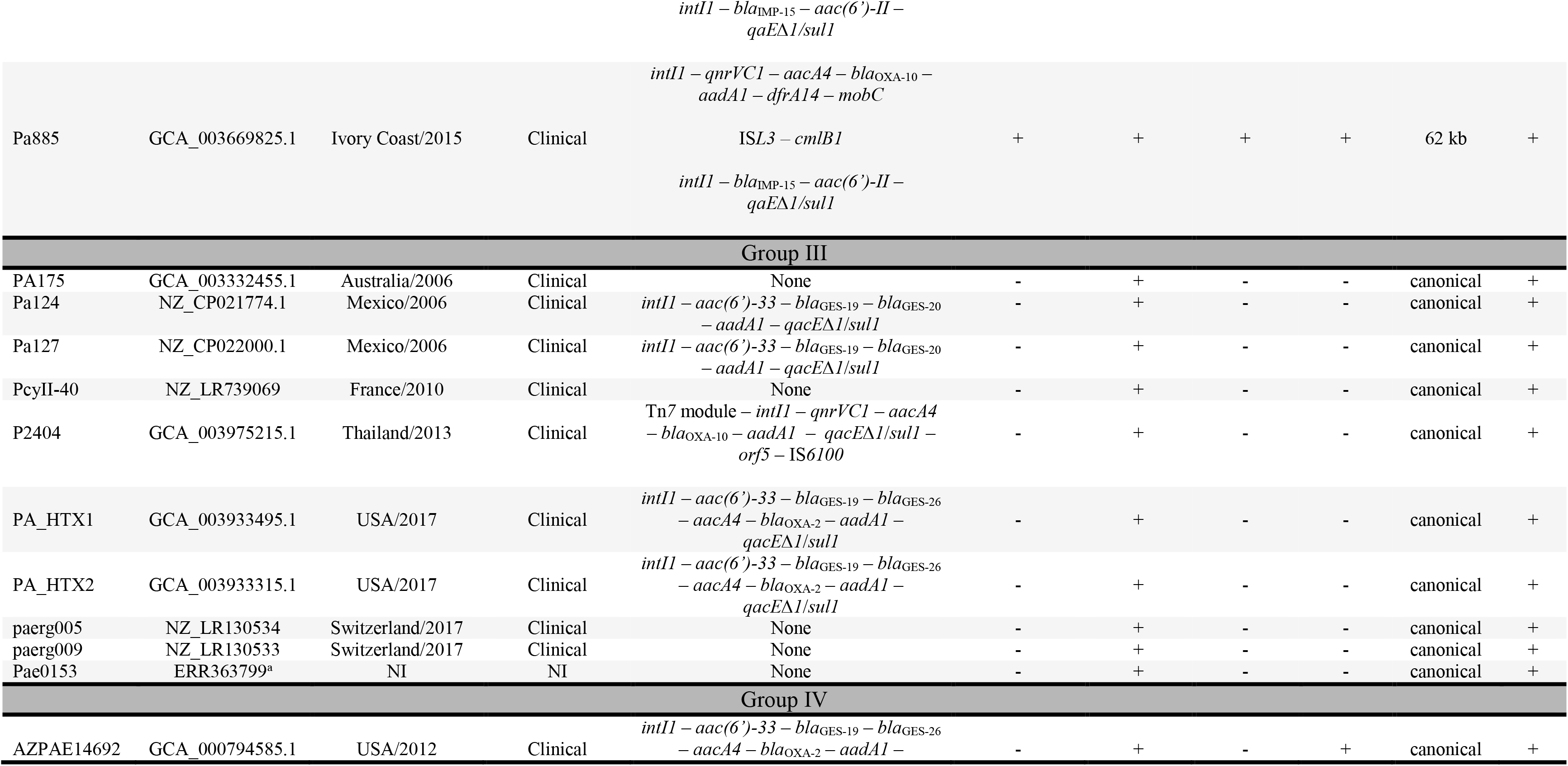
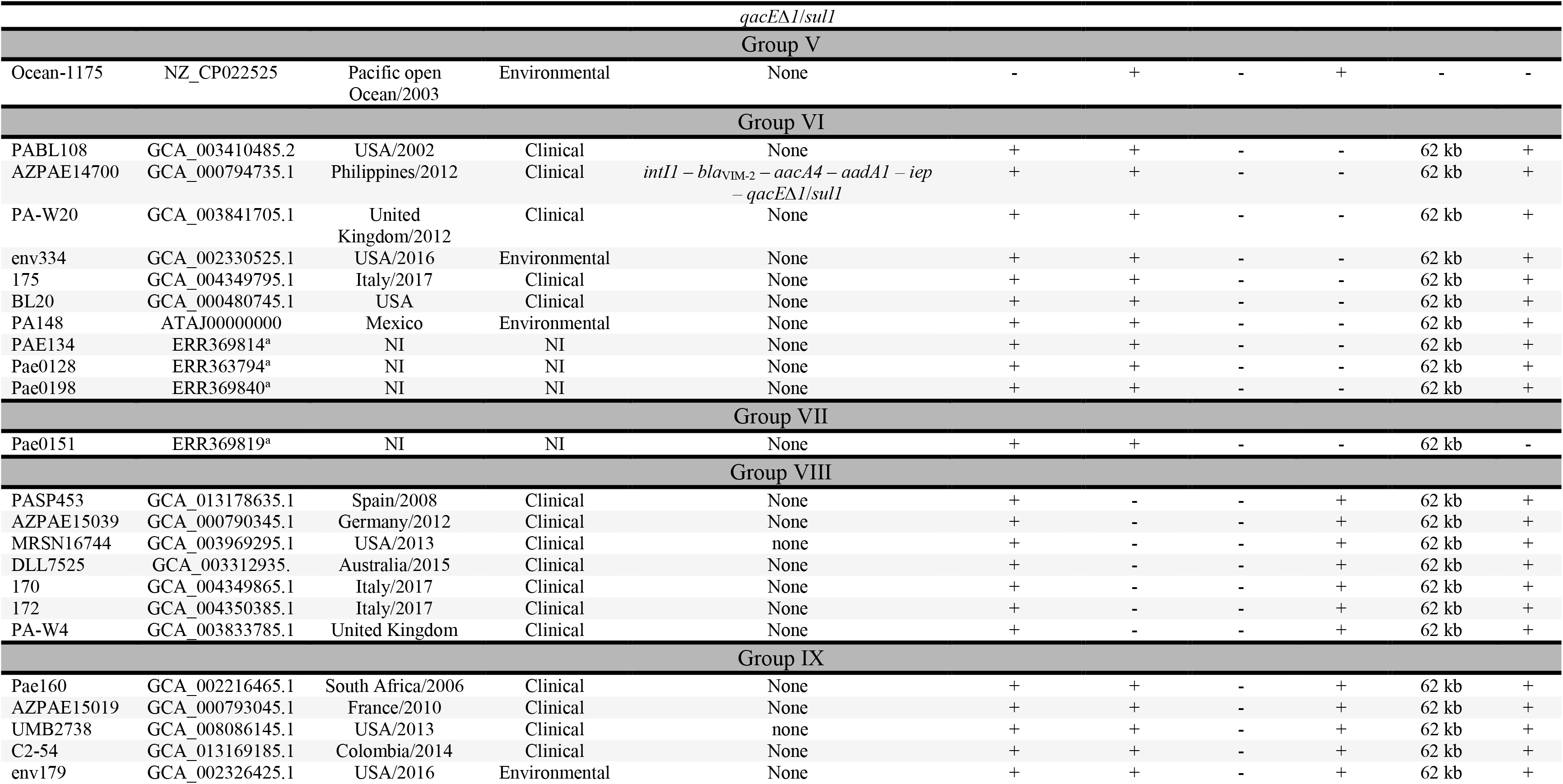
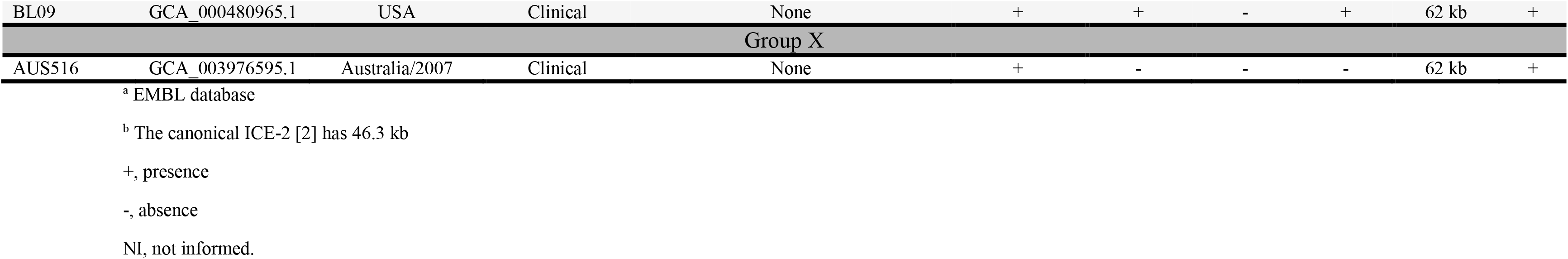
Epidemiological and genomic features of ST309 genomes analysed in this study.

ARG prediction was conducted in all ST309 genomes using the Comprehensive Antibiotic Resistance Database (CARD) [13]. The Resistance Gene Identifier (RGI) tool was used for predicting the resistome based on homology and SNP models with the default parameters.

Genomic island (GI) prediction was performed using the IslandViewer web server (http://www.pathogenomics.sfu.ca/islandviewer/) [14] and GIPSy software (https://www.bioinformatics.org/groups/?group_id1180). Integrative and Conjugative elements (ICEs) searches were performed using the ICEberg v2.0 platform [15].

The mobile genetic elements and virulence genes identified in PA834 genome were compared and mapped against the publicly available ST309 by performing BLASTn on NCBI’s nucleotide collection (nr/nt) database to verify intra-clonal variations concerning their accessory genomes.

### 2.4. Phylogenomic analysis

A phylogenomic reconstruction was performed using the 41 ST309 and 54 *P. aeruginosa* genomes from several epidemic/pandemic high-risk lineages. An orthology analysis using Roary pipeline [16] recovered 2,757 orthologous genes representing the core genome. These orthologues were concatenated (2.4 Mb) and submitted to the SNP-sites v.2.3.3 Program [17], which extracted 83 kb SNPs. A phylogenomic reconstruction was obtained, based on the concatenated SNPs, using the PhyML V3.1 [18] with GTR (general time reversible) substitution model, 100 bootstrap replications and the Maximum Likelihood method. All alignments were performed by MAFFT v7.271 [19], and the SNP tree was generated with iTOL v.3 [20].

## 3. Results and Discussion

### 3.1. ST309 is a pandemic lineage: emergence of a XDR strain in South America/Brazil

The PA834 strain was recovered from a fatal case of bloodstream infection that occurred in a hospital placed in the Brazilian Amazon Basin (Boa Vista city). The phenotypic analyses revealed that PA834 was resistant to a significant panel of antibiotics, which included 48 antimicrobial agents from more than 10 classes. It was only susceptible to aztreonam and polymyxins (MIC, 2 μg/mL for both polymyxin B and colistin), presenting, therefore, an extensively drug resistance profile (XDR) [11]. Moreover, the Hodge Test revealed that PA834 was a carbapenemase producer.

The MLST analysis revealed that PA834 belongs to ST309, which was recently considered a potential emerging threat, since it had been identified in Europe, Asia, North and Central America [5–9]. Moreover, we performed metadata analyses based on information recovered from BIGSDb database, GenBank and literature, and demonstrated that ST309 displayed a larger spatio-temporal distribution than previously reported. Our detailed analyses revealed that the ST309 has occurred in Oceania (Australia/2007-2015), Asia (Phillipines/2012, Korea/2014, Malaysia/2016, Thailand/2013), Africa (South Africa/2006, Ivory Coast/2012-2015), Europe (Greece, Spain/2008, Germany/2012, France/2007, 2010, 2011-2014, United Kingdom/2012, Italy/2017 and Switzerland/2017), North America (USA/2002, 2012 - 2017), Central America (Mexico/2006 and 2013) and South America (Colombia/2008 and Brazil/2016 – this study) (Table 1). Therefore, it is possible to assigned the ST309 as a pandemic lineage.

These genomes/strains were from both clinical and environmental sources, this latter including water, open ocean, dental care unit waterlines and hospital surfaces [5–9,21 and BIGSdb metadata]. Interestingly, ST309 has been continuoulsy isolated since the beginning of 2000 (Table 1) and, in spite of its huge distribution, no outbreaks had been reported, so far, to this lineage.

### 3.2. Phylogenomics of ST309

A phylogenomic reconstruction based on the core genome polymorphisms of ST309 genomes (n=42) and other epidemic/pandemic high-risk *P. aeruginosa* clones (n=54) was obtained. It was revealed that all ST309 genomes, recovered throughout 15 years from 7 continents (Table 1), grouped in a well defined cluster. No correlation between genetic relatedness and spatio-temporal occurrence was observed (Figure 1).

**Figure 1.**
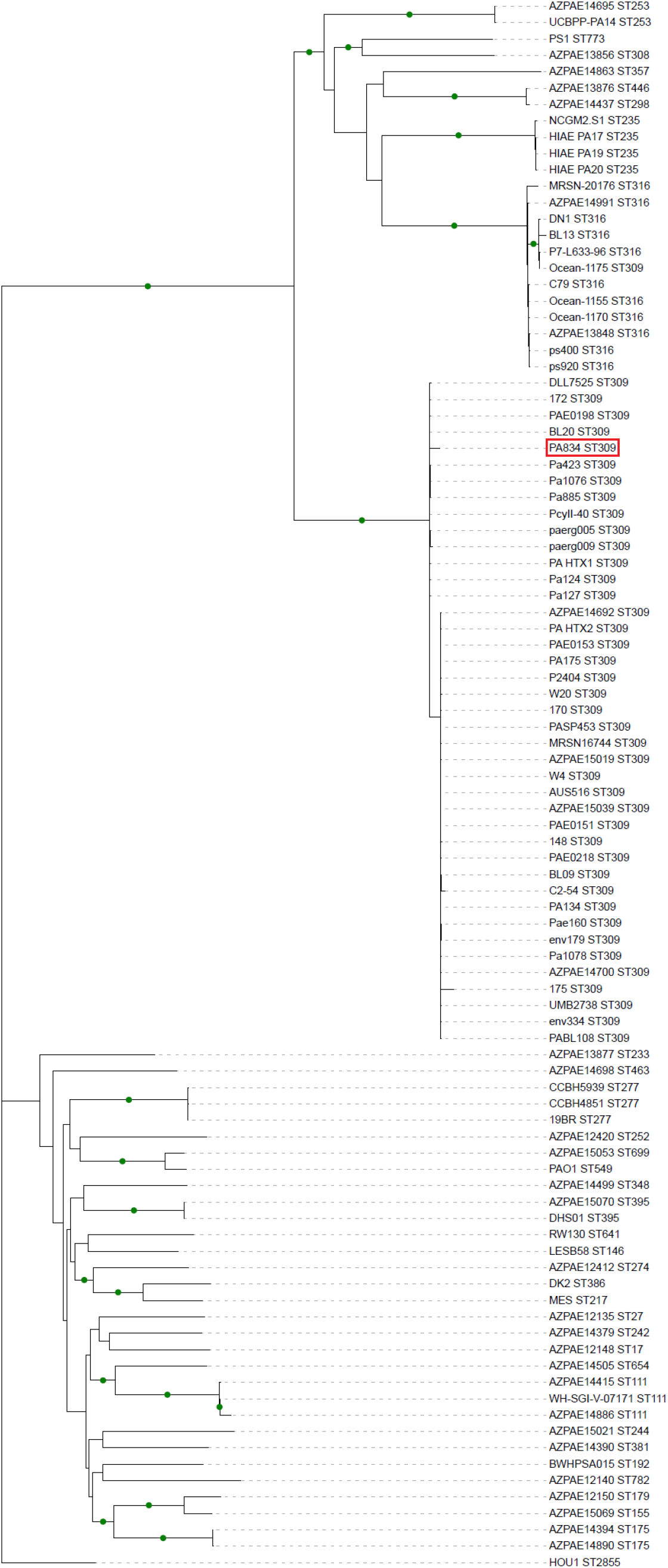
SNP-based phylogenomic tree of ST309 and other *P. aeruginosa* high-risk pandemic lineages. Topology based on bootstrap values >50 where green circles highlight bootstraps with values greater than 70. The name and corresponded ST of publicly available *P. aeruginosa* genomes from distinct high-risk pandemic lineages used in this reconstruction are indicated. The Brazilian ST309 genome PA834 obtained in this study is indicated by a red box.

### 3.3. The acquired resistome of ST309 lineage

The global resistome of ST309 lineage was composed by genes conferring resistance to β-lactams (*bla*_OXA-10_, *bla*_OXA-2_, *bla*_GES-19_, *bla*_GES-20_, *bla*_GES-26_), aminoglycosides (*aadA1, aacA4, aac(6’)-II, aac(6’)-33*), chloramphenicol (*cmlB1*), sulfonamides (*sul1*), ammonium quaternary compounds (*qacE∆1*), trimethoprim (*dfrA14*), carbapenems (*bla*_VIM-2_, *bla*_IMP-15_) and fluoroquinolones (*qnrVC1*). The most prevalent ARGs were *aadA1* and *sul1*, while *bla*_GES-20_, *bla*_VIM-2_ and *bla*_IMP-15_ were the rarest (Figura 2). In fact, *bla*_VIM-2_ and *bla*_IMP-15_, metallo-β-lactamase genes considered of clinical relevance, were only found in the genomes from the Brazilian PA834 strain and AZPAE14700 (*bla*_VIM-2_), and from Ivory Coast (*bla*_IMP-15_) (Table 1). Indeed, PA834 presented an imipinem MIC of >32 μg/mL besides a positive Hodge Test. Moreover, the higher number of ARGs were observed among the Ivory Coast genomes (Table 1), which included the *qnrVC1*. This gene was firstly identified in a clinical *Vibrio cholerae* strain recovered in the Amazon region in 1998 [22], however, *anrVCl* is considered rare in *P. aeruginosa*.

**Figure 2.**
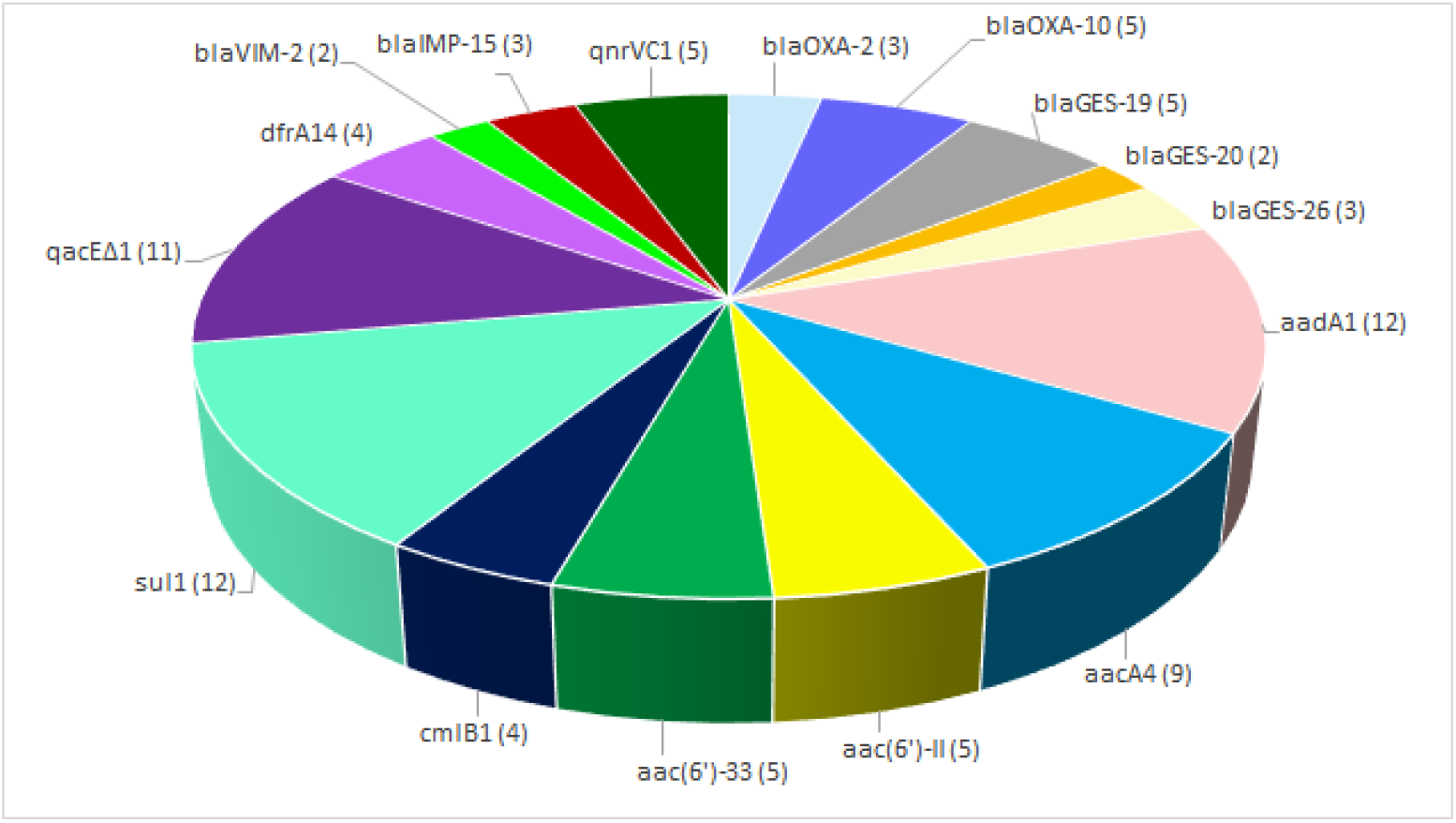
Distribution of ST309 acquired resistome. The ARGs identified among ST309 genomes are indicated together with the respective amount between parentheses.

Interestingly, 28/42 genomes had no acquired ARGs (Table 1), suggesting that the resistance phenotype of some strains from ST309 lineage was mainly due to intrinsic mechanisms. An example is the PA834 strain, which was resistant to 48 antibiotics, including all antipseudomonal drugs, but carried only a few ARGs involved with carbapenems, β-lactams, aminoglycosides and sulfonamide resistance (Table 1). This same pattern was observed in another high-risk pandemic *P. aerugnosa* lineage (ST395), in which intrinsic resistance mechanisms were the main responsible for the observed resistance phenotype [2].

#### 3.3.1. ARGs are associated to class 1 integrons

The resistome of ST309 was harboured by class 1 integrons, the exception was *cmlB1*, which was found in the context of the IS*L3* in Ivory Coast genomes (Table 1). Different from most of the ST309 genomes, which presented one class 1 integron, it was observed that PA834 and all Ivory Coast genomes harboured two integrons carrying ARGs (Table 1). One of these elements present in the Ivory Coast genomes (*intI1* – *qnrVC1* – *aacA4* – *bla*_OXA-10_ – *aadA1* – *dfrA14*) was also found in the unique genome from Thailand (Pa2404). In the case of PA834, besides the class 1 integron harbouring *bla*_VIM-2_ (Table 1), the other integron, which carried the *aac(6’)-II*, was contiguous to a Type 1 Restriction Modification System (*hsdSRM*). This system is involved with the attack and degradation of foreign DNA, as well as epigenetic gene regulation that influences the bacterial virulence and adaptation [23]. Global analysis demonstrated that this segment (85 – 95% coverage) was found in only three other genomes belonging to ST234 and ST773 pandemic lineages.

Our analyses revealed that the same integron array previously reported in PA_HTX1 and PA_HTX2 (USA, 2017) [5], was also present in AZPAE14692 (USA, 2012) (Table 1). Interestingly, the integron cassette arrays observed in these USA genomes and in Pa124 and Pa127 (Mexico, 2006) seemed to be the same element that evolved due to polymorphisms, such as the non-synonymous point mutation (Gly165Ser) that differs *bla*_GES-26_ from *bla*_GES-20_, and to acquisition/loss/reshufling of ARG cassettes (Figure 3).

**Figure 3.**
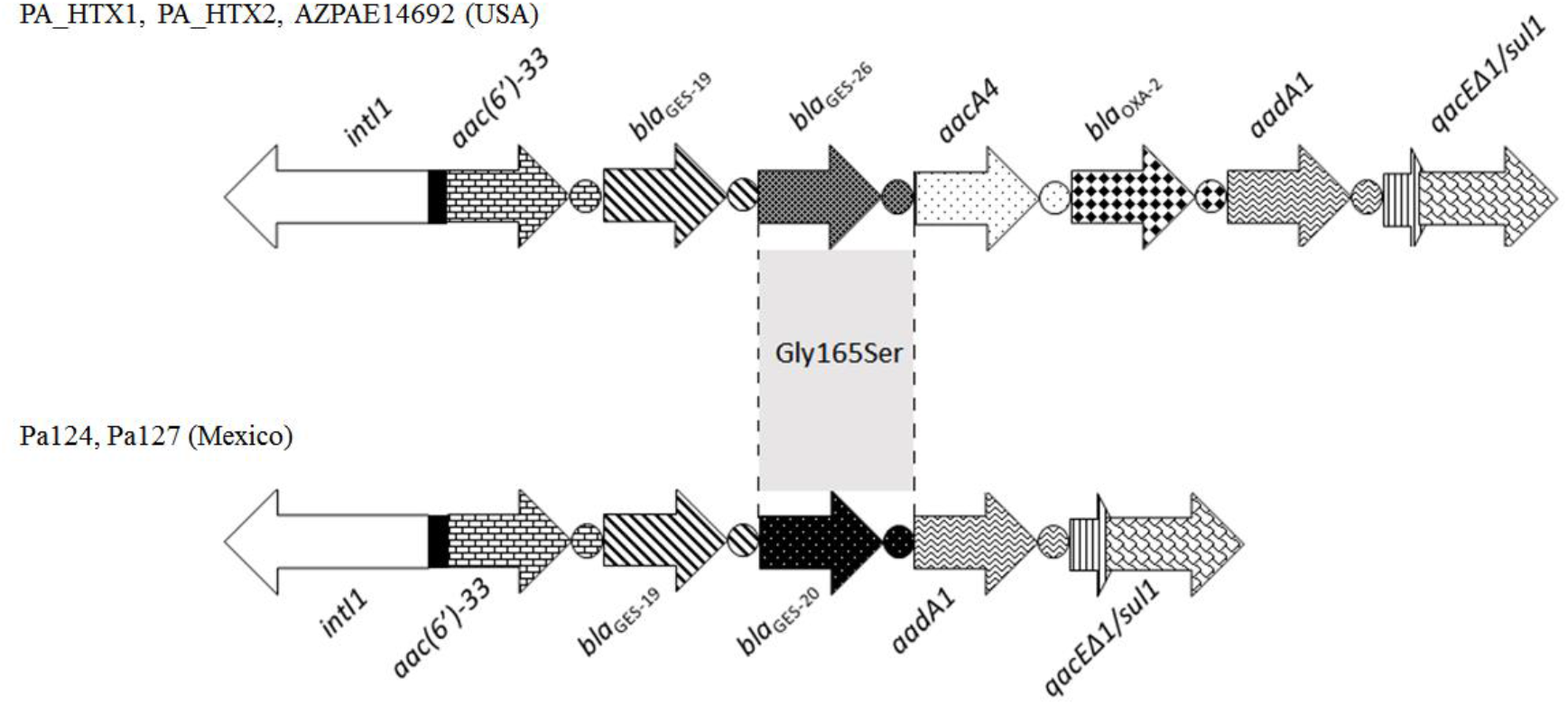
Schematic representation of the evolution of a class 1 integron circulating among ST309 genomes. Gene cassettes and their corresponding *attC* sites are represented by arrows and closed circles, respectively. The unique amino acid difference between *bla*_GES-20_ and *bla*_GES-26_ is indicated.

### 3.4. The mobilome of the ST309 lineage

Here it was revealed that the Brazilian PA834 and the other ST309 genomes harboured several mobile genetic elements including transposons, phages, ICE and GIs. As observed in Table 1, in most of cases there is no correlation between genomes sharing the same mobilome with their spatio-temporal distribution. In order to make this information clearer, the ST309 genomes in Table 1 was arranged in groups (I – X) according to the mobilome profile they shared.

#### 3.4.1. Genetic elements associated with heavy metal resistance

##### 3.4.1.1. The copper/silver/zinc-associated GI-7

A GI of 36.9 kb harbouring genes conferring heavy metal resistance was identified in 40/42 ST309 genomes (Figure 4a; Table 1). BLASTn analysis revealed that this island corresponded to GI-7 previously identified in strains of the high-risk ST308 and ST395 lineages [2,4]. ST308 lineage was found persistently colonizing the plumbing system of a healthcare unit in France [4], while the ST395 has persisted for 10 years in French hospitals, caused outbreaks in UK [2], and was also recovered from a metal contamined estuary [3]. Jeanvoine and Colleagues (2019) experimentally demonstrated that GI-7 had a crucial role in bacterium survival at copper contaminated environments, due to the presence of a set of genes conferring copper, silver and zinc resistance (*czcABC, copG, copM, copA, copB, copZ, copR, copS*, copper ABC transporter, plastocyanins) [4] (Figure 4a). This island shared 98% and 72% identity with sequences from the environmental species *Pseudomonas putida* and the phytopathogenic species *Pseudomonas syringae*, respectively. This suggests that GI-7 was probably originated from soil- and plant-associated environmental *Pseudomonas* and spread and established among high-risk clones of *P. aeruginosa*, such as ST308, ST395, ST111, ST253 [4], and now, ST309. Therefore, the presence of GI-7 in almost all spatio-temporally distributed ST309 genomes, indicated the role of this island in the success and establisment of ST309 as a pandemic lineage.

**Figure 4.**
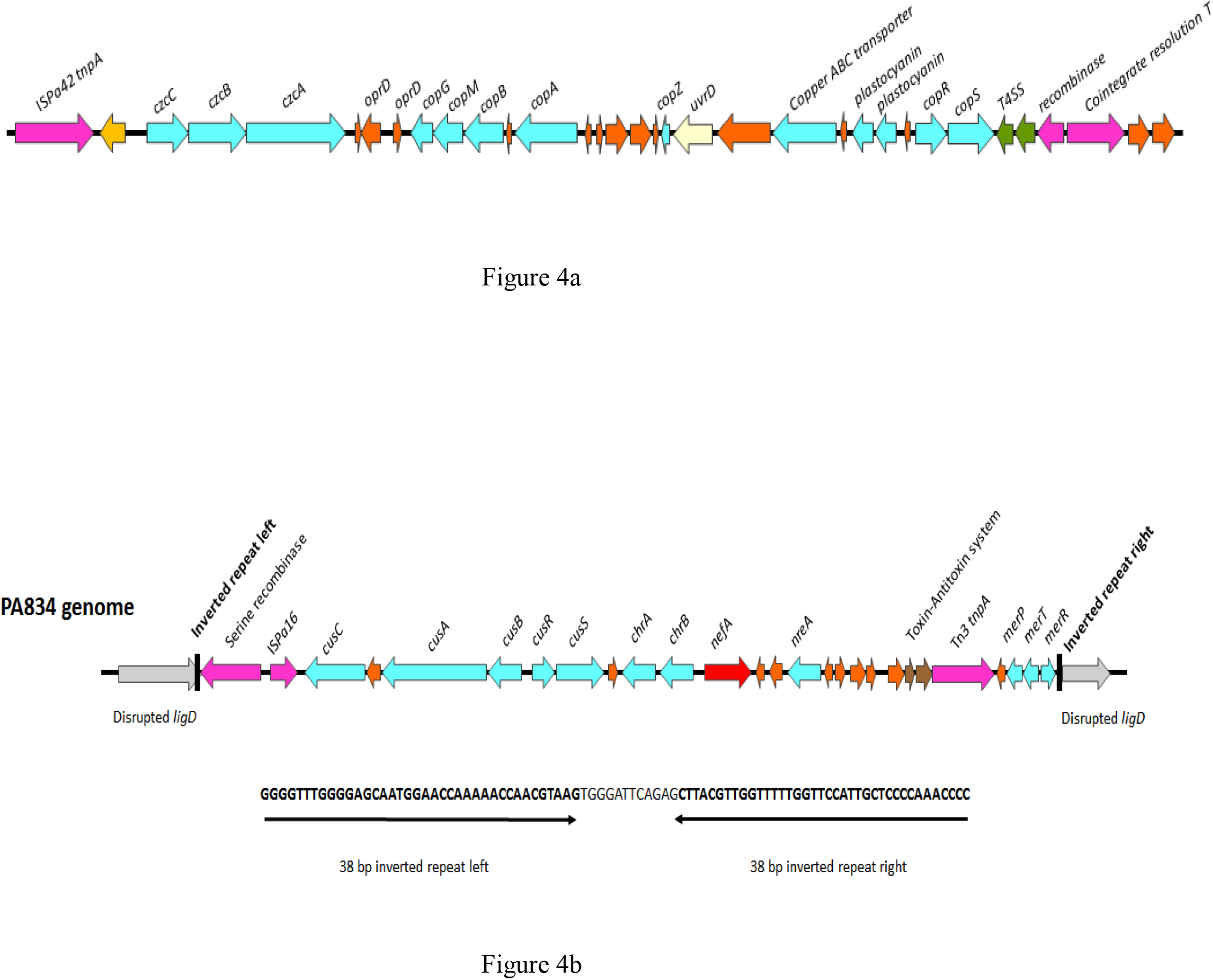
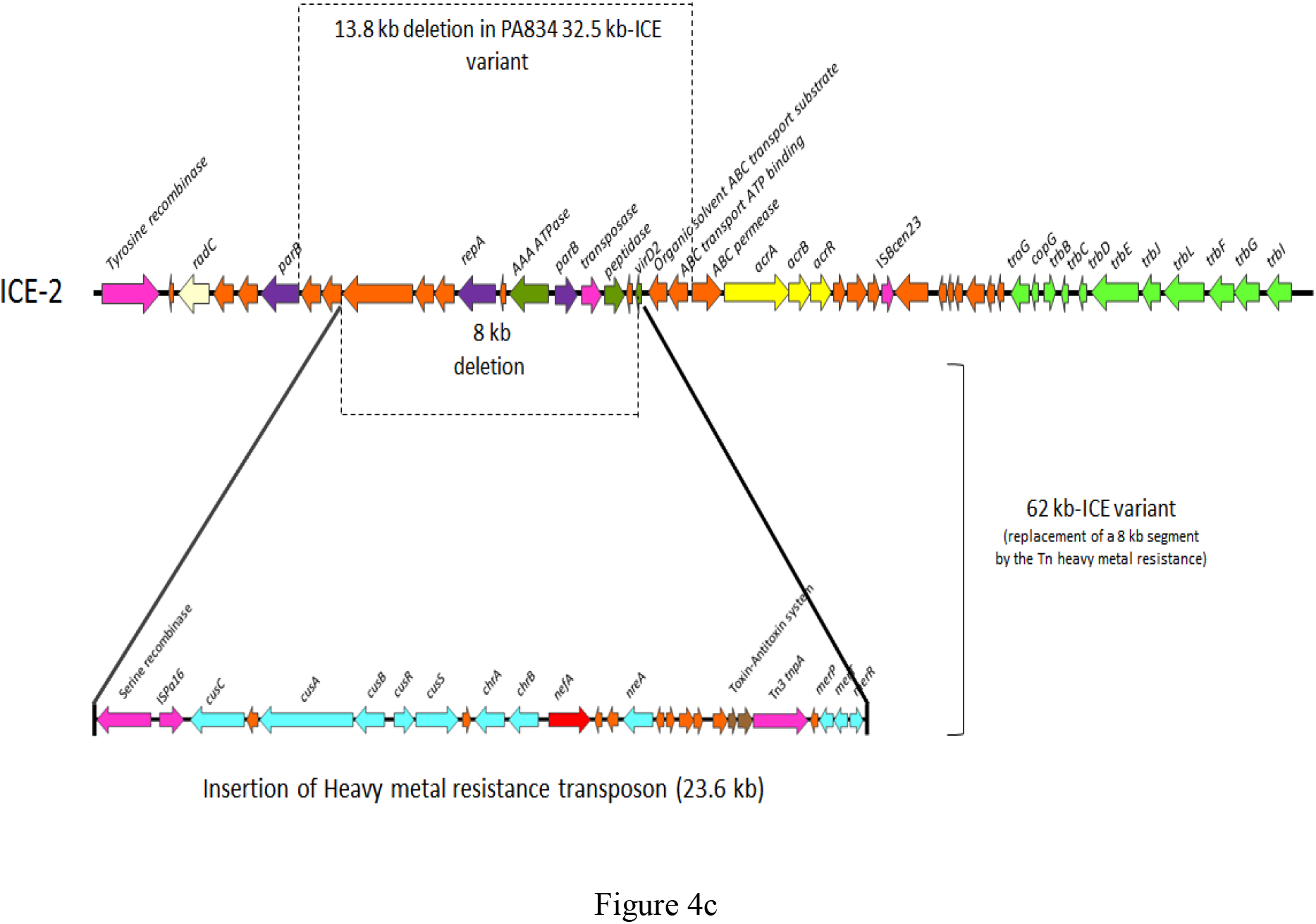
Schematic representation of ST309 mobilome harbouring heavy metal resistance determinants. (a) Structure of the heavy metal resistance genomic island GI-7 [4]. (b) Structure of the heavy metal resistance transposon inserted in PA834 chromosome. The inverted repeats sequence and the chromosomal location of transposon insertion are indicated. (c) Structure of ICE-2 and its variants found in ST309 genomes. The corresponded 13.8 kb deleted region of the 32.8 kb variant exclusively found in PA834 is indicated by dotted lines on the top of ICE-2 scheme. The events that led to the emergence of the 62 kb variant are indicated in the bottom of ICE-2 scheme: a 8 kb region was deleted (dotted lines) and replaced by the insertion of the heavy metal resistance transposon (solid lines) (for reference, see Figure 1b). In all schemes the genes are represented by arrows labelled with colours according to their main biological roles and functions: pink, recombinases; red, antibiotic resistance; turquoise blue, heavy metal resistance; light brown, Toxin-Antitoxin system; light green, Type IV Secretion System conjugal-associated genes; yellow, RND efflux pump; purple, replication; green, virulence-associated genes; nude, DNA repair; orange, hypothetical and other genes.

##### 3.4.1.2. Mercury resistance (mer) operon-harbouring element

The entire operon *merRTPADE* (mercury resistance) associated with the Tn*As2* from the Tn*3* transposon family was identified in 32/42 ST309 genomes. The *mer* operon is widespread among *Pseudomonas* in association with several elements. However, the genomic context of the *mer* operon observed here (IS*Pa38* – Tn*As2* – *merRTPADE*), was exclusively found in ST309 genomes and as part of the ICE-5 identified in ST395 strains from clinical origin (DHS01) and metal contamined water (E67) [2,3]. One environmental ST309 genome recovered from Pacific Open Ocean in 2003 (Ocean-1175) harboured the mercury resistance operon in a distinct mobile genetic element, which also included the arsenic resistance operon *ars* (*merRTPCADE* - IS*30 tnpA* – *arsBADCR* – recombinase gene – Tn*SA3*). Interestingly, this same element was found in genomes from ST316 (Ocean-1170 and Ocean-1155) also recovered from Pacific Ocean in 2003. Considering the presence of this element shared between ST309 and ST316 recovered from the same niche suggests the occurrence of lateral transfer among bacteria even in vast and dynamics environments such as oceans [24].

##### 3.4.1.3. Heavy metal resistance transposon

A 23.6 kb transposon associated with IS*Pa16*, belonging to the Tn*5* family, carrying a remarkable arsenal of heavy metal resistance genes was found disrupting the *ligD* gene in PA834 genome (Figure 4b). Interestingly, this transposon was also found in 29/41 publicly available ST309 genomes, however, they were embedded in another genomic context different from *ligD*, as discussed below (subsection 3.4.1.4). This element harboured additional copies of *merP*, *merT* and *merR*; the *nreA* and *nreB* genes, involved with nickel resistance; the *chrA* and *chrB* genes, related with chromate resistance; the gene cluster composed by the two component system *cusSR* and the *cusBAC*, responsible for copper and silver resistance. BLASTn analysis revealed that this transposon (95-100% coverage and 98-100% identity) was found in genomes and plasmids from several environmental bacterial species, many of them inhabiting contamined niches: a *P. aeruginosa* recovered from an oil contaminated soil in Kwait in 2012 (Genbank accession no. CP025229.1); a nitrogen-fixing and rhizosphere-associated *P. stutzeri* (Genbank accession no. CP002622.1); a *P. balearica* recovered from corrosion products from a corroded pipe in Australia in 2016 (GenBank accession no. CP045858.1); a *P. putida* involved with mercury methylation recovered from soil in China in 2014 (GenBank accession number CP022560.1); a *P. soli* found in a wastewater in Korea (GenBank accession no. CP009365.1); and a plasmid from an freshwater algae-associated *Stenotrophomonas rhizophila* recovered in China in 2016. Therefore, these findings are evidences that this heavy metal resistance transposon, similarly to GI-7, probably contributes to bacterial survival in stressful conditions such as heavy metal contamined sites.

##### 3.4.1.4. The antiseptic/heavy metal resistance-associated ICE-2 and its variants exclusively found in ST309

The mobilome mining and comparative genomic analyses revealed that 11 ST309 genomes harboured the ICE-2 (46.3kb) previously found in DHS01, a clinical strain from the high-risk ST395 recovered from France in 1997 [2] (Figure 4c). Genomic BLASTn revealed that, besides ST395, the ICE-2 was present in other *P. aeruginosa* high-risk pandemic lineages, such as ST235, ST111, ST175 as well as in local STs, such as the XDR ST260 associated with severe infection cases [25,26]. Except by Ocean-1175, the remaining ST309 genomes harboured variants of ICE-2 that arose due to indels of gene blocks. A smaller variant of 32.5 kb, which suffered a 13.8 kb deletion considering the canonical 46.3-kb ICE-2 sequence, was exclusively found in the Brazilian PA834 genome (Figure 4c). A larger version of ICE-2 (62 kb) was found in the 29 remaining ST309 genomes (Figure 4c). It arose as a consequence of a 8-kb deletion considering the canonical 46.3 kb ICE-2, which was replaced by the insertion of a 23.6 kb segment. Interestingly, this segment corresponded to the aforementioned heavy metal resistance transposon that was found disrupting the chromosomal *ligD* gene in PA834 (subsection 3.4.1.3). Although the 23.6 kb heavy metal resistance transposon had already been identified in several environmental species, as discussed above (subsection 3.4.1.3), the entire 62-kb ICE variant seems to be, so far, unique to these ST309 genomes. It is worth to mention that the observed gain and loss of large DNA segments in both ICEs of 32.5 kb (deletion of 13.8 kb) and 62 kb (replacement of a 8kb DNA segment by the heavy metal resistance transposon of 23.6 kb) occurred in the same region of ICE-2, indicating that this region represents a hotspot in this element.

All ICE variants identified here carried an homologue of *acrRAB-tolC* efflux pump from the RND superfamily. This efflux pump is essential for bacterial survival and colonization/virulence, especially during the course of infection when the pathogen contacts toxic substances or adheres with the host cells [27]. Additionaly, it has an important role in resistance to several antibiotics (macrolides, fluoroquinolones, β-lactams, tetracyclines, chloramphenicol, rifampin, novobiocin, fusidic acid), basic dyes (acriflavine and ethidium) and detergents (bilesalts, Triton X-100, SDS) [28]. Moreover, a recent study has also demonstrated the role of this efflux pump in the carbapenem resistance in *Escherichia coli* [29].

It is of note that a remarkable arsenal of heavy metal resistance genes harboured by mobile genetic elements was unraveled in ST309. As aforementioned, several of these elements were also found in other high-risk lineages, particularly in ST395 recovered from both clinical and heavy metal contamined environments, which shared with ST309 the GI-7, ICE-2 and part of ICE-5 [2–4]. Interestingly, strains from ST395 and ST309 have already been identified sharing the same niche [30], where horizontal gene transfer events could have take place. However, the circulation of these genetic elements among different pandemic lineages (ST309, ST395, ST308, ST111, ST253…) suggests that a common *P. aeruginosa* ancestor had horizontally acquired such elements from an environmental source or that they emerged in environmental *Pseudomonas* species before its transfer to high-risk lineages [2,4].

#### 3.4.2. Genetic elements associated with virulence

The *pilLNOPQRSUVM* gene cluster coding for the type IV thin pili, involved with plasmid conjugation, twitching motility, biofilm formation and adherence onto surfaces such as catheters [31,32], was present in PA834. This *pil* gene cluster was found in the context of the IS*66* Family, and it was identical to that found in the pKLC102 ICE from the epidemic *P. aeruginosa* clone C that affected cystic fibrosis patients in Europe [33]. Twenty of 41 ST309 genomes harboured the segment containing the *pil* genes sharing 95% - 100% nucleotide identity with that identified in PA834 (Table 1).

Interestingly, the *zona ocludens* (*zot*) toxin gene, related to disassembly of the intercellular tight junctions of mammalian cells and invasiveness, was exclusevely found in PA834 genome. This genome had two *zot* genes, one harboured by *ISPa32*, together with T2SS genes, two copies of the antitoxin *hipB* and several prophage genes. The other copy was carried by the IS*Pa1328*, which also harboured genes coding for phage proteins and for ParE/ParD toxin/antitoxin system, involved with plasmid maintenance.

## 4. Conclusion

This study reported the emergence of ST309 in South America/Brazil, providing insights about its global epidemiology, which allowed the assignment of ST309 as a pandemic high-risk lineage. The unraveled ST309 mobilome and resistome revealed the presence of several mobile elements harbouring a remarkable amount of heavy metal resistance genes, many of them shared with other high-risk *P. aeruginosa* lineages. These elements, together with the identified ARGs and virulence determinants, probably contributed to ST309 adaptive fitness, which had been crucial for its evolution, persistence, spread and successful establishment as a pandemic lineage.

## Ethical Approval

This study was approved by the participating Ethics Committee of the Federal University of Roraima (CEP-UFRR) under the number CAAE 54446116.2.0000.5302. All methods were performed in accordance with the relevant guidelines and regulations.

## Author contributions

EF – Conceptualization and design of the study, performed the experiments, analyzed and interpreted the data, drafted, reviewed and edited the manuscript; FF and SM – performed the experiments; RC – collected the bacterial strains; ACV - Conceptualization and design of the study, scientific supervision, funding acquisition, revision, edition and final approve the manuscript. All authors have read and agreed to the published version of the manuscript.

